# MIG-21 is a novel regulator of Wnt and Netrin signaling in gonad migration identified from published scRNA-seq data and functionally validated in *C. elegans*

**DOI:** 10.1101/2025.02.24.639896

**Authors:** Xin Li, Kacy Lynn Gordon

## Abstract

Using a recently published scRNA-seq dataset of adult *C. elegans* hermaphrodites, we identified a previously unknown regulator of the germ line stem cell niche (the distal tip cell, or DTC). The gene *mig-21* has the highest “marker score”–yet no known role–in the DTC. Using classical genetics techniques, RNAi knockdown, and live cell imaging, we discovered that *mig-21* integrates information from the Wnt and Netrin pathways to guide anteroposterior and dorsoventral DTC migration. Our study demonstrates the utility of scRNA-seq datasets in revealing testable hypotheses about genetic networks that were masked by redundancy in traditional screening methods.

## Introduction

Single-cell RNA sequencing (scRNA-seq) approaches chemically label cDNA made from transcripts from single cells before sequencing and then identify cell types by transcriptome similarity via clustering algorithm (Zhang et al. 2023). scRNA-seq has rocketed to prominence in studies of developmental biology in the past decade (Svensson et al. 2020). Cell atlases constructed by single-cell RNA-sequencing are being generated for model organisms and human organs across a range of genetic and physiological states, enabling for example the discovery of rare cell populations with importance in human disease (Plasschaert et al. 2018). Ideally, scRNA-seq datasets will lead to new biology being explored well beyond their initial publications. Like genome sequences, these datasets could become precious tools not only for further bioinformatic analyses, but for hypothesis generation for functional genetics studies.

We used a recently published *C. elegans* scRNA-seq dataset (Ghaddar et al., 2023) to study a cell of interest: the *C. elegans* hermaphrodite gonad stem cell niche, called the distal tip cell (DTC). Each of the two gonad arms of the *C. elegans* hermaphrodite has a single DTC. The DTC is a migratory germline stem cell niche that has long served as a model for cell migration and stem cell niche biology (Hubbard and Greenstein 2000; Gordon 2020). Its stereotyped migration during post-embryonic development patterns the correct U-shaped gonadal morphology of each adult gonad arm. In the L2 and early L3 larval stages, the DTCs migrate away from each other along the ventral body wall in an anterior or posterior direction. In the L3 larval stage, each DTC makes a 90-degree turn off the ventral body wall and then another 90-degree turn onto the dorsal body wall to face the midbody. These turns pattern the bend region of the mature gonad. During the L4 larval stage, each DTC migrates along the dorsal body wall and comes to rest at the dorsal midbody (Hubbard and Greenstein 2000).

The genetics of DTC migration and cessation have been studied for decades. DTC migration is governed by two major signaling gradients: a ventral-to-dorsal UNC-6/Netrin gradient that primarily governs D/V migration (the first turn) and a posterior-to-anterior EGL-20/Wnt gradient that primarily governs A/P migration (the second turn) (Wong and Schwarzbauer 2012). Proper A/P and D/V guidance of the DTC results from the integration of inputs from both networks, and they demonstrate some degree of redundancy (Levy-Strumpf & Culotti, 2014). During reproductive adulthood, the DTC is stationary, highly elaborated, and continues to signal to the underlying germ stem cells to maintain them in an undifferentiated state. We asked if previously unknown regulators of the DTC could be identified by examining highly expressed genes detected in that cell type by scRNA-seq.

We used the web app WormSeq.org (https://wormseq.org) for the dataset reported by (Ghaddar et al., 2023) to look for DTC “marker genes”. Surprisingly, the gene with the highest “marker score” is *mig-21*, which has been studied in the context of neuronal cell migration but was not known to function in the DTC. *mig-21* was initially identified in a screen for genes that affect touch receptor neuron development (Du and Chalfie 2001), many of which proved to be essential for proper anterior-posterior migration of the progenitors of these cells. MIG-21 is a thrombospondin repeat (TSPI)-containing transmembrane protein that interacts with the Netrin receptor UNC-40/DCC to regulate cell polarization and migration of Q neuroblasts during early larval development (Middelkoop et al. 2012). Like the two DTCs, the two Q neuroblasts initially migrate away from one another along the A/P axis (Sundararajan and Lundquist 2012). MIG-21-dependent polarization of QL and QR restricts the threshold response to the EGL-20/Wnt gradient in these cells (Middelkoop et al. 2012). The role of *mig-21* in Q neuroblast migration along the same Wnt and Netrin signaling gradients that guide DTC migration makes it a plausible candidate to examine in the DTC.

In this study, we established a role for *mig-21* in DTC migration, focusing on the Wnt and Netrin signaling pathways that regulate both Q neuroblast and DTC migration. We also examined whether known interactors of *mig-21* in Q neuroblasts coregulate DTC migration. Finally, we examined whether a known regulator of DTC migration cessation interacts with *mig-21*. This work provides new insights into the complex interplay of signaling pathways that guide DTC migration and highlights the important role of *mig-21*, emphasizing the power of scRNA-seq in hypothesis generation for probing gene function in development.

## Results and Discussion

### DTC migration is regulated by *mig-21*

We identified *mig-21* as our gene of interest by looking for DTC-expressed genes with a high “marker score” (genes with relatively high expression levels and relatively specific expression) and “specificity score” (determined by Jensen-Shannon distance)(https://cole-trapnell-lab.github.io/monocle3) in scRNAseq results (Ghaddar et al., 2023). The gene *mig-21* is at the top of the “marker” list and is third for “specificity” in the DTC (Figure 1A). It is the only gene that appears in the top ten genes of both sorts.

**Figure 1.**
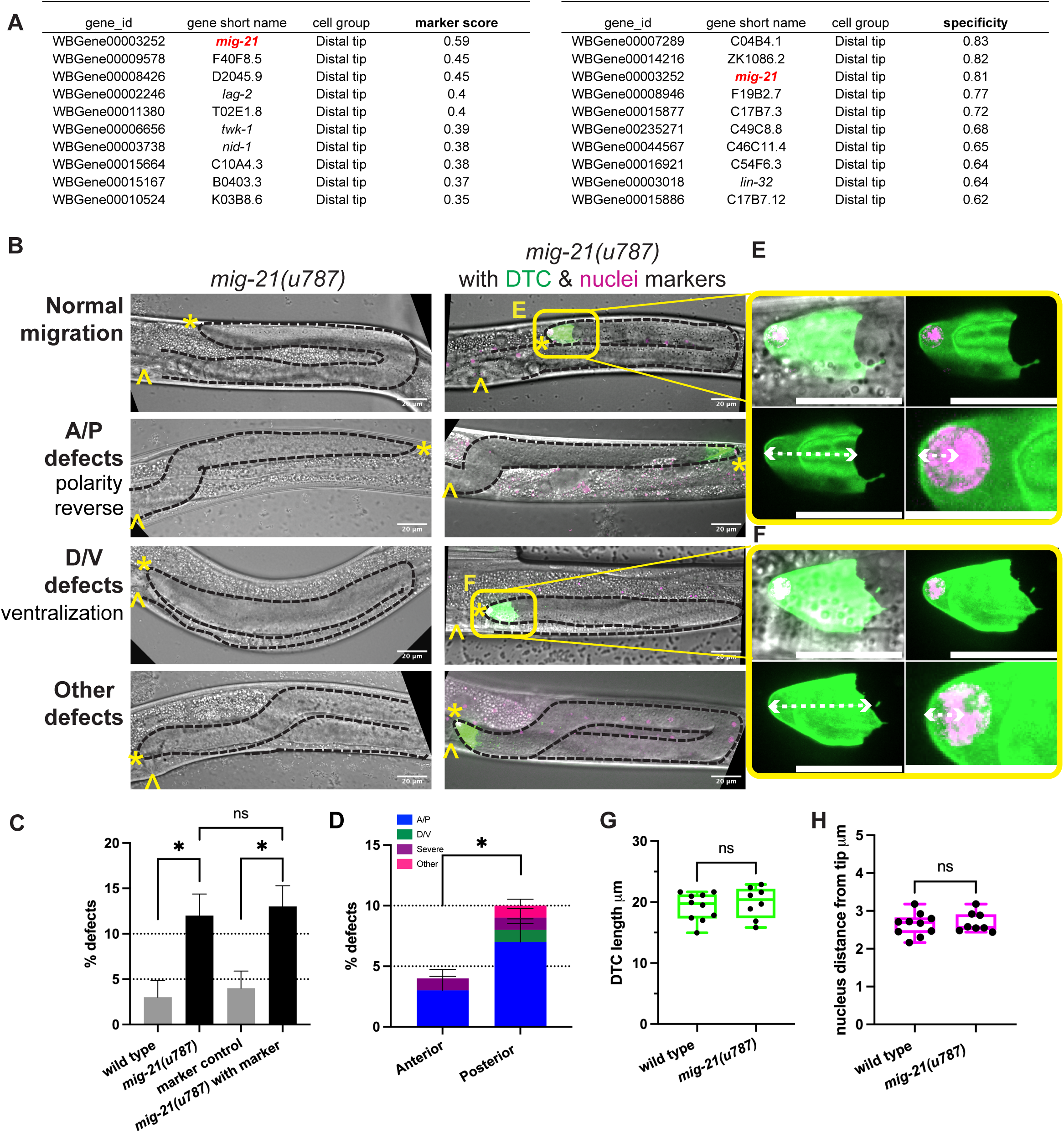
*mig-21* regulates DTC migration but not cell structure (A) Top 10 genes ranked by marker score (left) and specificity (right) from WormSeq.org (https://wormseq.org) for the distal tip cell at the young adult stage. *mig-21* is highlighted as the only gene on both lists. (B) Micrographs on the left: DIC imaging of *C. elegans* hermaphrodites at the late larval L4 stage for *mig-21(u787)* worms. Micrographs on the right: DIC merged with fluorescence imaging of the same stage *mig-21(u787)* crossed with a strain bearing a transgene that marks the membrane of the DTC, *cpIs122[lag-2p::mNeonGreen:: PLC^δPH^*], and a nuclear marker inserted at the endogenous *lag-2* locus *lag-2(bmd202[lag-2::P2A::H2B::mT2])* (Singh et al. 2024). Images are Z-projections through the thickness of the gonad required to capture the whole distal gonad. Black dashed lines outline gonads. Anterior left and ventral down. Yellow asterisks mark DTC; yellow carets mark the proximal vulval position. Scale bar: 20 μm. Yellow boxes indicate the positions of insets shown in E and F. (C) Percentage of DTC migration defects across experimental groups. Low-penetrance but significant defects were identified in *mig-21(u787)* alone and with fluorescence markers. Wild type N2 strain control (n=3/92), *mig-21(u787)* (n=23/189), p < 0.05; markers control (n=4/104), *mig-21(u787)* with markers (n=28/215), p < 0.05. Introducing fluorescence markers did not significantly alter *mig-21(u787)* defect rates (p > 0.05). (D) Percentage of worms with migration defects in anterior vs. posterior gonad arms of *mig-21(u787)* worms. Migration defects were more frequent in the posterior gonad arm (n=18/189) compared to the anterior arm (n=7/189), p < 0.05. Anterior-posterior polarity (A/P) defects (n=19/189) were significantly more common than dorsal-ventral polarity (D/V) defects (n=1/189), p < 0.0001. (C-D) All sample sizes refer to individual worms. Error bars represent the standard error of the sample proportion. Statistical analysis was performed using a pairwise proportion test, with p-values adjusted for multiple comparisons via the Benjamini-Hochberg procedure. Significant differences are indicated between groups where applicable. ****p < 0.0001 ***p < 0.001 **p < 0.01; *p < 0.05; ns=no significant. The corresponding sample sizes and statistics are presented in Table S1. (E-F) Enlargement of distal tip fluorescence images from 1B, showing normal (E) and defective (F) migration. Annotations show measurements quantifying DTC morphology and nuclear localization (graphed in 1G-H). White dashed lines represent measurement parameters: (Bottom panels Left) DTC length, as the linear distance from the distal tip to the proximal boundary of the DTC; (Bottom panels Right) nuclear position, as the distance from the distal tip to the geometric center of the nucleus. Scale bars 20 μm except bottom right panels 10 μm. (G) Box plots overlaid with all datapoints measuring the DTC length. Sample sizes refer to individual gonads. Wild type N2 control n=10, and *mig-21(u787)* n=8. Statistical significance was calculated by unpaired, two-tailed Student’s t-tests, error bars represent ±SEM. No significant difference was observed, t=0.6080, df=16, 95% confidence interval −1.737 to 3.135 μm, p = 0.7374. (H) Box plots overlaid with all datapoints measuring the DTC nuclei position. Sample sizes refer to individual gonads. Wild type N2 control n=10 and *mig-21(u787)* n=8. Statistical significance was calculated by unpaired, two-tailed Student’s t-tests, error bars represent ±SEM. No significant difference was observed, t=0.4361, df=16, 95% confidence interval −0.2280 to 0.3461 μm, p = 0.7531.

First, we examined the effect of *mig-21* loss-of-function on DTC migration by analyzing a strain bearing the putative null *mig-21(u787)* mutant allele (Middelkoop et al. 2012) which has a premature amber stop codon in the extracellular domain. Amber stop codons are not read-through in *C. elegans* (Patel et al. 2008). In *mig-21(u787)* mutants, we observed a low penetrance defect in DTC migration in which some DTCs failed to execute the two turns correctly (Figure 1B-C).

We categorized defects into anterior/posterior (A/P) vs. dorsal/ventral (D/V) polarity defects according to the specific phase of migration that was affected, as well as “other” migration defects and “severe” defects in which gonad growth failed or formed a disorganized mass (Fig. 1B and Fig. S1A). *mig-21(u787)* mutants exhibited a significantly higher proportion of A/P defects than D/V defects, with more defects observed in the posterior gonad arm than in the anterior (Figure 1D).

DTC migration defects are caused by signaling defects (Levy-Strumpf and Culotti 2014), as well as by cell structural abnormalities including those that arise from defects in Rac GTPase signaling (Singh et al., 2024; Reddien & Horvitz, 2000) or actomyosin-based contractility and nuclear mispositioning (Agarwal et al. 2024). To further investigate the potential role of *mig-21* in influencing DTC shape or nuclear position during migration, we crossed *mig-21* mutants with a DTC marker strain that has membrane-localized DTC fluorescence and a nuclear histone tag *(lag-2p::mNeonGreen::PLC^δPH^; lag-2(bmd202[lag-2::P2A::H2B::mT2])* (Singh et al. 2024). The presence of these transgenes did not enhance DTC migration defects in either a wild-type or *mig-21(u787)* background (Figure 1C). Defects caused by knockdown of Rac GTPase genes were not significantly enhanced by *mig-21(u787)* (Fig. S1B-D).

In the L4 stage, both control and *mig-21(u787)* mutant DTCs maintained their normal shape and displayed well-defined nuclei polarized in the direction of migration (Figure 1E-H), suggesting that *mig-21* likely interacts with signaling pathways involved in DTC guidance moreso than cytoskeletal dynamics or nuclear positioning, with a specific impact on anterior-posterior directional polarity, especially in the posterior gonad arm.

### *mig-21* regulates Wnt signaling in the DTC

Wnt signaling is the main regulator of DTC and Q neuroblast migration along the A/P axis, the latter dependent on *mig-21* (Sundararajan & Lundquist, 2012). Redundancy among Wnt pathway members and between Wnt and Netrin signaling had previously been described by revealing epistasis tests (Levy-Strumpf & Culotti, 2014; Levy-Strumpf et al., 2015). Genetic redundancy revealed by superadditive or synergistic interaction does not suggest true molecular redundancy between Wnt and Neterin pathway members, but redundancy in the guidance information they impact. We hypothesized that *mig-21* may also function redundantly with Wnt signaling to guide A/P migration, explaining why it had not previously been discovered as a regulator of DTC migration.

To assess the functional interaction between *mig-21* and the Wnt signaling pathway, RNAi by feeding was used to knock down Wnt pathway genes in the *mig-21(u787)* background. We targeted *mom-5* (which encodes the main Frizzled receptor that acts during DTC migration to guide A/P polarity) (Levy-Strumpf et al., 2015), *lin-17* (another Frizzled) (Levy-Strumpf & Culotti, 2014), and five Wnt ligand genes, *egl-20* (which forms a Wnt gradient crucial for Q neuroblast migration (Pani & Goldstein, 2018), *lin-44*, *mom-2, cwn-1, and cwn-2*. We predicted that if *mig-21* acts redundantly to the Wnt signaling pathway, we will see an enhancement of Wnt knockdown phenotypes in *mig-21(u787)* mutants. After empty-vector control RNAi treatment, *mig-21(u787)* mutants showed the same minor ∼10% A/P migration defect rate we observed on standard growth media.

Knockdown of *mom-5*/Frizzled with RNAi in wild-type worms caused a ∼20% total per worm A/P migration defect (Fig. 2A-B), which agrees with previous *mom-5* RNAi results (Levy-Strumpf et al., 2015; Singh et al., 2024). In the *mig-21(u787)* background, the incidence of A/P migration defects after *mom-5* RNAi increases to over 50% (Fig. 2A-B), with an almost complete bias for the posterior DTC (Fig. 2C). Incidence of A/P migration defects caused by *lin-17* RNAi treatment of wild-type worms was more modest (<10%), but was also enhanced over twofold by *mig-21(u787)* with the opposite bias, for the anterior DTC (Fig. 2C). We thus conclude that *mig-21* functions redundantly with Wnt receptors during DTC migration, playing a bigger role as a counterpart to *mom-5* in the posterior DTC and a minor role with *lin-17* in the anterior DTC. Because the *mig-21(u787)* DTC migration defect is so modest, the strong synergistic effect of the double loss of function leads us to conclude that *mig-21* and Frizzled genes function redundantly at the genetic level. At the pathway level, we do not propose that MIG-21 may function truly redundantly as a Wnt receptor, but potentially that redundancy among Wnt pathway members themselves depends on *mig-21* function.

**Figure 2.**
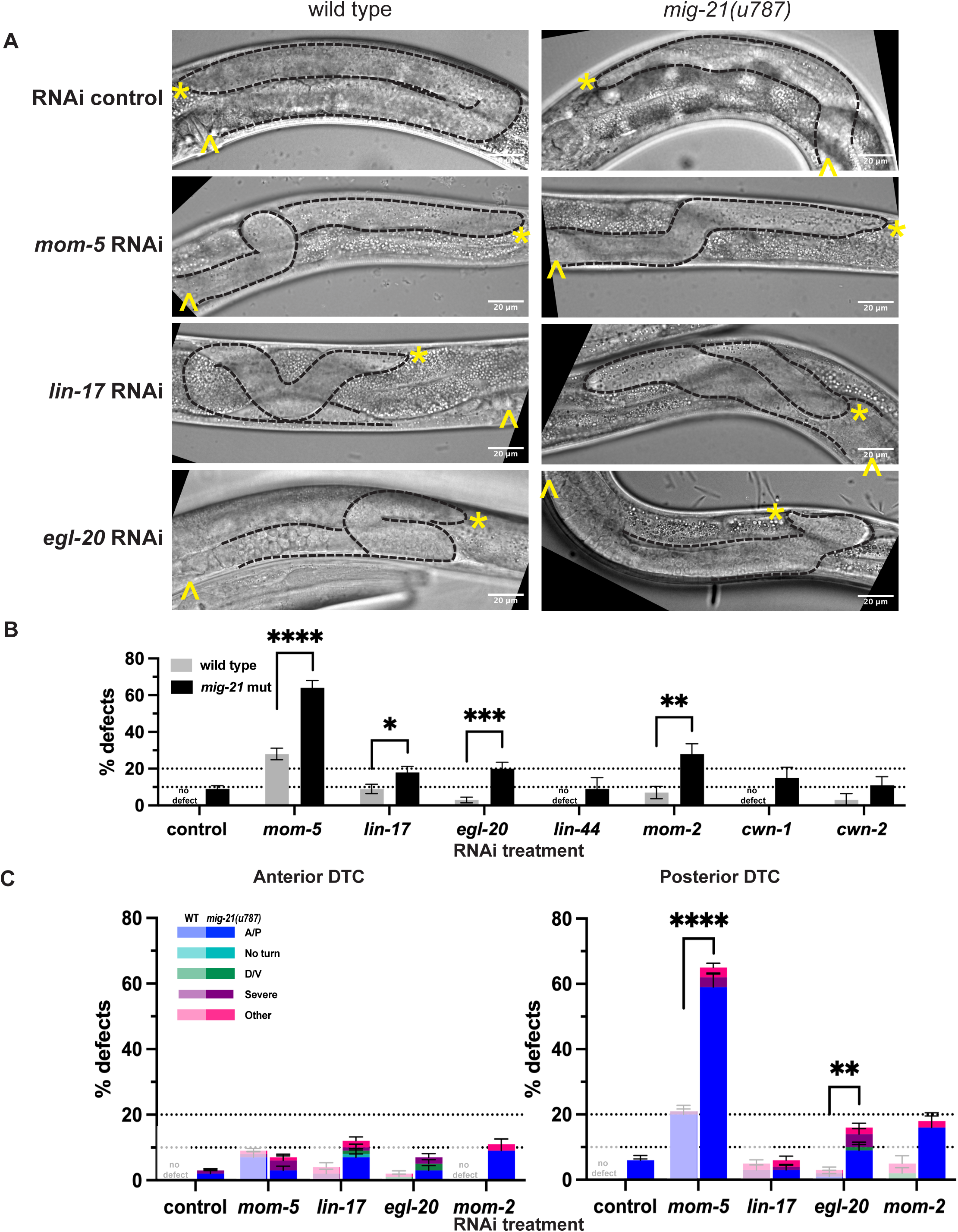
*mig-21* regulates Wnt signaling in the DTC (A) Micrographs: DIC imaging of *C. elegans* hermaphrodites at the late larval L4 stage, comparing wild type N2 (left) and *mig-21(u787)* (right) under RNAi control L4440 empty vector, *mom-5* RNAi, *lin-17* RNAi, or *egl-20* RNAi feeding treatment to assess DTC migration defect phenotypes. Images are Z-projections through 2-3 μm showing the distal gonad. Anterior left and ventral down. Black dashed lines outline gonads. Yellow asterisks mark DTC; yellow carets mark the proximal vulval position. Scale bar: 20 μm. (B) All DTC migration defects across experimental groups, including wild type N2 (gray) and *mig-21(u787)* (black) strains, under RNAi control L4440 empty vector, Frizzled receptors *mom-5* and *lin-17* and Wnt ligands *egl-20, lin-44, mom-2, cwn-1, cwn-2* RNAi feeding treatments. Significant enhancements of migration defects by *mig-21(u787)* were observed in *mom-5*, *lin-17*, *egl-20*, and *mom-2* groups. (C) DTC migration defects for the groups with significant enhancement by *mig-21(u787)* in anterior (left) and posterior (right) arms. Lighter one means wild type groups, darker one means *mig-21(u787)* groups. (B-C) All sample sizes refer to individual worms. No defect means no defect observed in that group. Error bars represent the standard error of the sample proportion. Statistical analysis was performed using a pairwise proportion test, with p-values adjusted for multiple comparisons via the Benjamini-Hochberg procedure. Significant differences are indicated between groups where applicable. ****p < 0.0001 ***p < 0.001 **p < 0.01; *p < 0.05; no mark means the comparison was not statistically significant. The corresponding sample sizes and statistics are presented in Table S2.

A Wnt ligand-independent function of *lin-17*/Frizzled has recently been discovered in the earliest asymmetries in the somatic gonad (So et al. 2024), however, DTC migration is Wnt-dependent (Levy-Strumpf & Culotti, 2014). We next tested genetic interactions of Wnt ligand genes with *mig-21*. When they have been investigated previously, a single loss of function of the five Wnt ligand genes causes negligible DTC migration defects (Levy-Strumpf & Culotti, 2014). Indeed, we did not observe major A/P migration defects after wild-type worms were exposed to RNAi for any of the Wnt ligand genes. The *mig-21(u787)* mutation significantly enhanced migration defects of *egl-20*/Wnt and *mom-2*/Wnt RNAi. We conclude that *mig-21* works partially redundantly with liganded Frizzled receptors (primarily MOM-5) to guide A/P polarity of (primarily posterior) DTC migration, with special sensitivity for EGL-20, the Wnt ligand to which *mig-21* regulates the response in the Q neuroblasts (Middelkoop et al. 2012).

### *mig-21* interactors in the Q neuroblasts also act in the DTC

In the Q neuroblasts, MIG-21 partners with PTP-3, which encodes a LAR receptor phosphotyrosine-phosphatase (RPTP) (Harrington et al. 2002), to regulate Wnt-responsive cell polarization. *ptp-3* is not known to function in the DTC, though its transcripts are detectable in the adult (https://wormseq.org) and L2 DTC (Cao et al. 2017). RNAi knockdown of *ptp-3* in wild-type worms caused minor incidence of gonad migration defects (<10%, Fig. 3A-B). However, *mig-21(u787)* mutants treated with *ptp-3* RNAi had a gonad defect rate of ∼20%, with the majority of defects being A/P migration defects in the posterior gonad (Fig. 3C).

**Figure 3.**
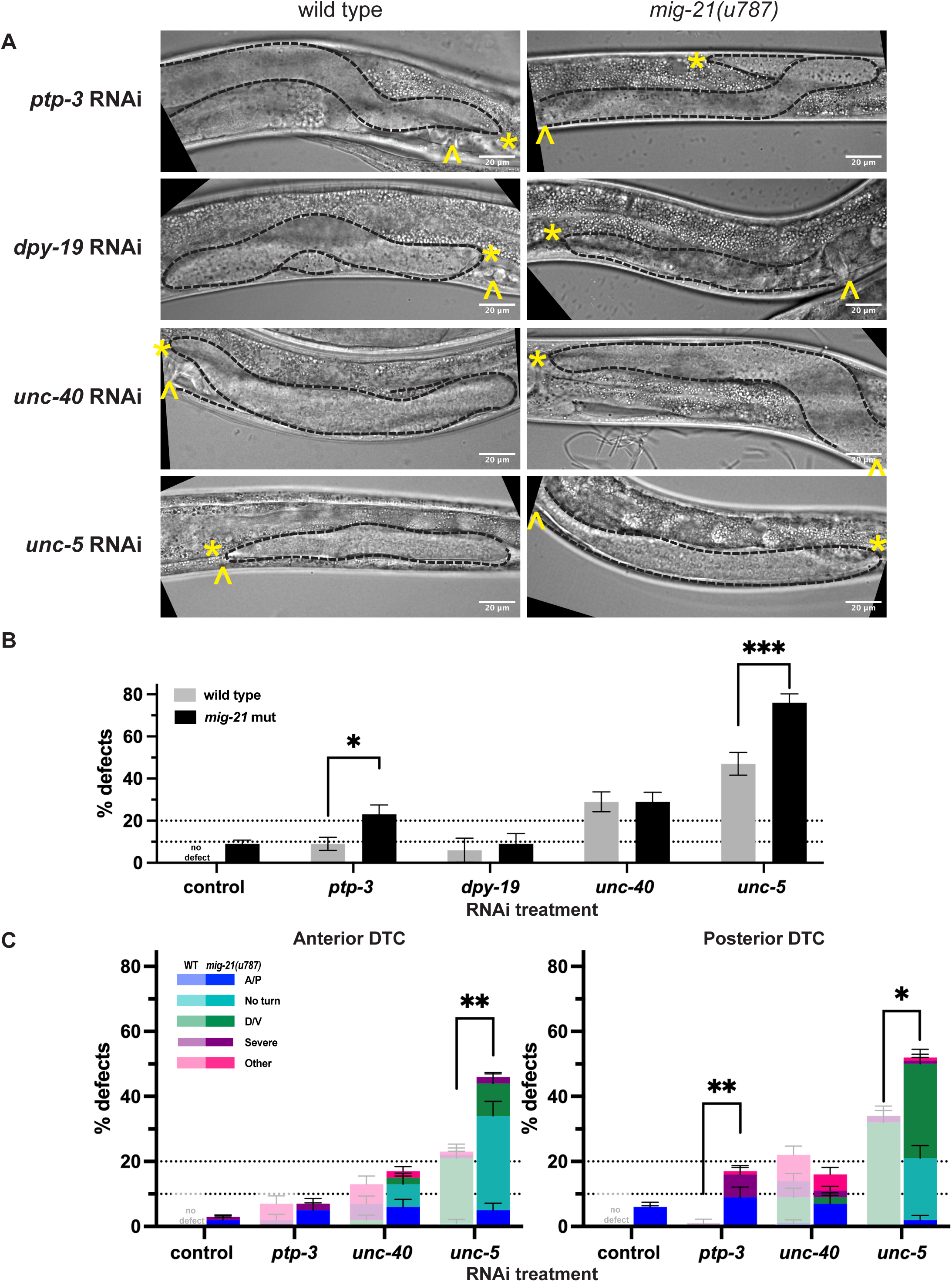
*mig-21* works with *ptp-3* and regulates Netrin signaling in the DTC (A) Micrographs: DIC imaging of *C. elegans* hermaphrodites at the late larval L4 stage, comparing wild type N2 (left) and *mig-21(u787)* (right) under RNAi control L4440 empty vector, LAR receptor *ptp-3*, C-mannosyltransferase *dpy-19* and Netrin receptors *unc-40, unc-5* RNAi feeding treatment. Images are Z-projections through 2-3 μm showing the distal gonad. Anterior left and ventral down. Black dashed lines outline gonads. Yellow asterisks mark DTC; yellow carets mark the proximal vulval position. Scale bar: 20 μm. (B) All DTC migration defects across experimental groups, including wild type N2 (gray) and *mig-21(u787)* (black) strains, under RNAi control L4440 empty vector. *ptp-3* RNAi, *dpy-19* RNAi, *unc-40* RNAi, or *unc-5* RNAi feeding treatment. Significant enhancements were observed in *ptp-3* (p < 0.05) and *unc-5* (p < 0.001) groups, but not *dpy-19* (p > 0.05) and *unc-40* (p > 0.05) group. (C) DTC migration defects across different experimental groups in anterior (left) and posterior (right) arms (exclude *dpy-19* group). Lighter one means wild type groups, darker one means *mig-21(u787)* groups. Significant enhancement was also observed in posterior arm of *ptp-3* (p < 0.01) group. Significant enhancement was observed in anterior (p < 0.01) and posterior (p < 0.05) arms of *unc-5* group, with a highly significant increase in the “no turn” phenotype was observed in both the anterior (p < 0.0001) and posterior (p < 0.0001) arms of *unc-5* group. While *mig-21(u787)* does not affect the incidence of gonad defects in the *unc-40* RNAi group, it does change the character of migration defects. (B-C) All sample sizes refer to individual worms. No defect means no defect observed in that group. Error bars represent the standard error of the sample proportion. Statistical analysis was performed using a pairwise proportion test, with p-values adjusted for multiple comparisons via the Benjamini-Hochberg procedure. Significant differences are indicated between groups where applicable. ****p < 0.0001 ***p < 0.001 **p < 0.01; *p < 0.05; ns = no significant. The corresponding sample sizes and statistics are presented in Table S3.

Several common factors both regulate the DTC and interact with LAR. The guanine nucleotide exchange factor Trio interacts with LAR in mammalian cells (Debant et al. 1996); in *C. elegans unc-73*/Trio loss of function phenocopies Rac GTPase loss of function in the DTC (Singh et al. 2024). Substrates of LAR, like Beta-catenin and cadherin, are known as regulators of the post-migratory DTC (Gordon et al., 2019; Tolkin et al., 2024). Finally, the LAR receptor mediates adhesion between germline stem cells and the *Drosophila* male germline stem cell niche (Srinivasan et al. 2012). Future work on the role of *mig-21/ptp-3* cooperation in the DTC is warranted.

Another interactor of *mig-21* in Q neuroblast migration, *dpy-19,* encodes a C-mannosyltransferase (Buettner et al. 2013). RNAi knockdown of the *dpy-19* gene alone or in combination with *mig-21(u787)* has the same minor DTC migration defect as *mig-21(u787)* alone, suggesting that any role of *dpy-19* in the DTC is likely limited to its processing of *mig-21* (Fig. 3A-B).

Finally, the Netrin receptor UNC-40/DCC interacts with MIG-21/PTP-3 negatively in QR and acts in parallel in QL. We find that wild-type worms on *unc-40* RNAi have a ∼30% overall DTC migration defect, including ventralization of migration (Fig. 3A-B). When *mig-21(u787)* mutants are put on *unc-40* RNAi, the overall migration defect penetrance does not change or differ between anterior and posterior DTCs, but the nature of the phenotypes changes to include failure to turn at all, and other A/P polarity defects in both the anterior and posterior DTC which are never observed after *unc-40* knockdown of wild-type worms, along with notable suppression of D/V polarity defects in the posterior DTC (Fig. 3C). Our results suggest that *mig-21* and *unc-40* have complex and potentially differing interactions in the two DTCs, as they do in the two Q neuroblasts. However, UNC-40 is not the only Netrin receptor in the DTC.

### *mig-21* regulates the Netrin pathway in the DTC

The Netrin signaling ligand UNC-6 establishes a concentration gradient along the dorsal-ventral (D/V) axis, directing cellular and growth cone migration (Norris & Lundquist, 2011). This guidance function is achieved through interaction with its receptors, UNC-40/DCC (Chan et al. 1996), and the Netrin receptor UNC-5 (Leung-Hagesteijn et al. 1992). Netrin signaling confers the dominant D/V polarity information in DTC migration, and UNC-5 regulates D/V DTC migration both independently and redundantly with UNC-40 in transducing a repulsive ventral UNC-6/Netrin signal (Hedgecock et al., 1990), and in a Netrin-independent manner (Levy-Strumpf et al., 2015). Since *mig-21* alters the nature of *unc-40* RNAi defects in the DTC, we hypothesized *mig-21* may interact with UNC-5/Netrin receptor during DTC migration as well.

Loss of *unc-5*/Netrin receptor function causes ventralization of DTC migration in which the DTC migrates out and back along the ventral body, never crossing to the dorsal body wall (Hedgecock et al., 1990). The *mig-21(u787)* allele alone rarely shows evidence of D/V migration defect (<5%, Fig. 3C). Treatment of wild-type worms with *unc-5* RNAi causes a ∼50% overall ventralization defect (Fig. 3A) (with a slight posterior bias, Fig. 3C), which is in line with previous observations (Levy-Strumpf & Culotti, 2014; Singh et al., 2024). Treatment of *mig-21(u787)* mutants with *unc-5* RNAi enhanced per-worm D/V migration defects to over 75% (∼50% each for anterior and posterior DTCs). Notably, over 20% of affected gonads in both the anterior and posterior failed to turn (Fig. 3C). The “no turn” phenotype is considered to result from defects of both A/P and D/V polarity of the DTC, with the DTC failing to cross to the dorsal side and then failing to turn back towards the midbody (Levy-Strumpf & Culotti, 2014). We thus conclude that *mig-21(u787)* enhances the frequency and severity of migration defects caused by the loss of function of UNC-5/Netrin receptor and likely acts redundantly in the same pathway, especially in its contribution to A/P polarity.

### *mig-21* mediates Wnt-Netrin pathway crosstalk

Netrin pathway signaling confers the dominant D/V polarity information, and Wnt pathway signaling confers the dominant A/P polarity information during DTC migration, however, the two signaling networks function somewhat redundantly at the genetic level (Levy-Strumpf & Culotti, 2014). Wnt and Netrin pathway loss of function can both mutually enhance and also suppress DTC migration defects caused by loss of function in the other pathway, revealing that each pathway contributes to both proper A/P and D/V DTC migration. Because of its apparent redundancy with both Wnt and Netrin receptor genes, we next wondered if *mig-21* might act as a relay or convergence point between the two signaling pathways.

We tested how the *mig-21(u787)* allele affected DTC migration in genetic contexts with combined Wnt and Netrin loss of function. We first generated an *mig-21(u787); unc-5(e152)* double mutant. The *unc-5(e152)* allele has a premature stop codon truncating the intracellular domain of all isoforms (Mahadik & Lundquist, 2023; Killeen et al., 2002; Hedgecock et al., 1990); that intracellular domain mediates the repulsion that brings about the first turn (Lee et al. 2005). The *unc-5(e152)* mutant and the double mutant have no appreciable A/P migration defect beyond that of *mig-21(u787)* alone (Fig. 4A-B). The *unc-5(e152)* mutant has an overall D/V migration defect of ∼70% (with a posterior bias, Fig. S2). This is a more penetrant defect than *unc-5* RNAi produces, and it is not enhanced by our *unc-5* RNAi (Fig. 4C), suggesting that the *unc-5(e152)* mutant is a complete loss of function for *unc-5*-mediated regulation of DTC migration. We find that this migration defect rate is not significantly enhanced by *mig-21(u787)* (Fig. 4B-C), providing further evidence that *unc-5* (via its C-terminal region) and *mig-21* act in the same DTC migration pathway and in the same direction.

**Figure 4.**
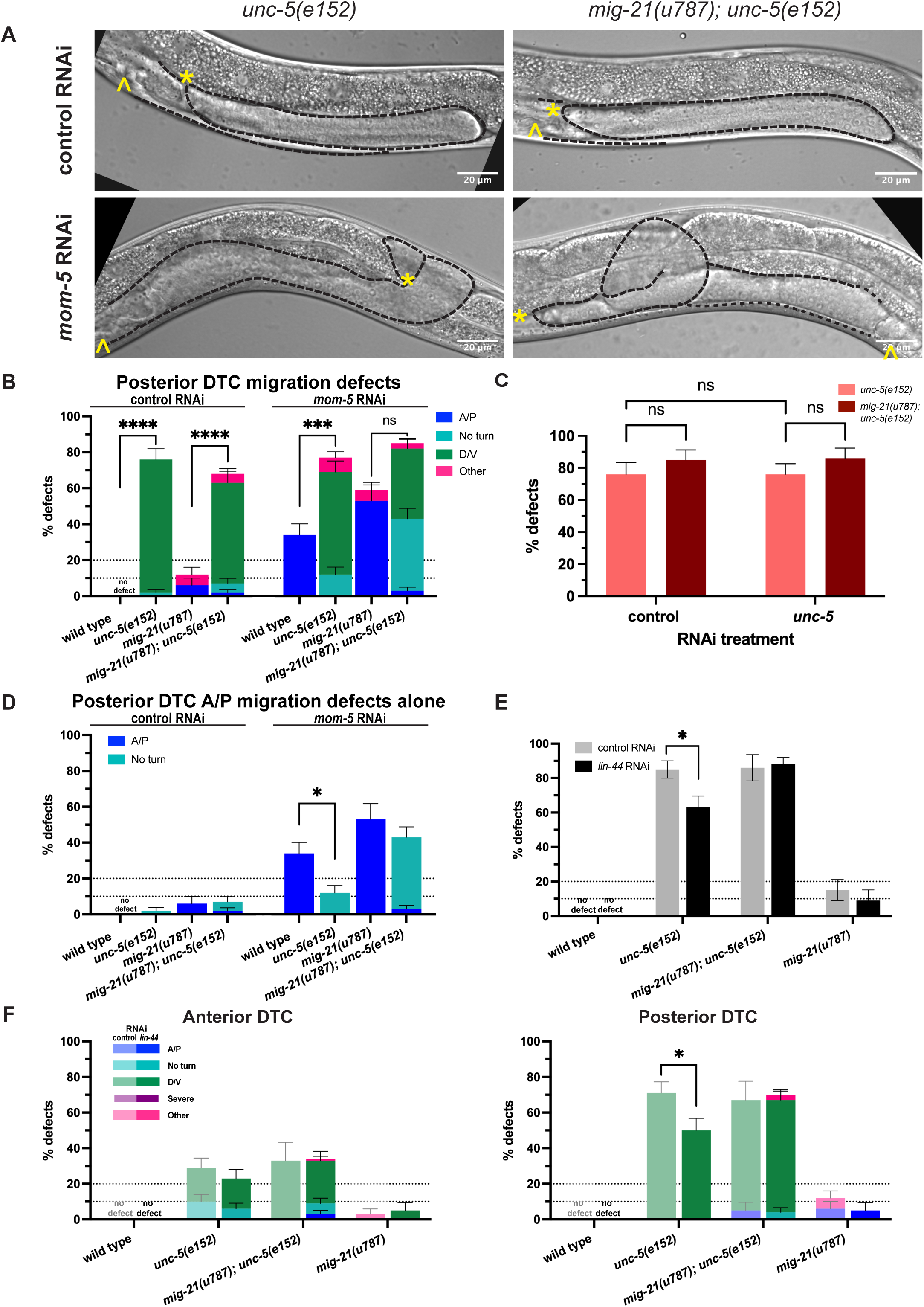
*mig-21* mediates Wnt-Netrin pathway crosstalk in the DTC (A) Micrographs: DIC imaging of *C. elegans* hermaphrodites at the late larval L4 stage, comparing single mutants *unc-5(e152)* (left) and double *mig-21(u787); unc-5(e152)* (right) mutants under RNAi control L4440 empty vector and *mom-5* RNAi feeding treatment to assess DTC migration defect phenotypes. Images are Z-projections through 2-3 μm showing the distal gonad. Anterior left and ventral down. Black dashed lines outline gonads. Yellow asterisks mark DTC; yellow carets mark the vulval position. Scale bar: 20 μm. (B) Comparing the percentage of all classes of migration defects observed across different experimental groups in posterior gonad arms only. (C) All DTC migration defects across experimental groups comparing *unc-5(e152)* (salmon) and *mig-21(u787);unc-5(e152)* (cayenne) strains, under control RNAi L4440 empty vector and *unc-5* RNAi feeding treatments. (D) Comparing the percentage of only the classes containing anterior-posterior migration defects in posterior gonad arms shown in 4B. Anterior-posterior migration defects include “A/P” polarity reverse and “no turn” categories. Isolating these defects from the “D/V” and “other” classes makes it easier to see the significant suppression of *mom-5-*RNAi-induced A/P migration defects which is lost in *mig-21(u787);unc-5(e152)* double mutants on *mom-5* RNAi. (E) Comparing the percentage of all the DTC migration defects observed across different experimental groups under *lin-44* RNAi feeding treatment. The rate of *unc-5(e152)* migration defects (n=44/52) was significantly suppressed by *lin-44* RNAi (n=34/54). However, this suppression is lost in the *mig-21(u787); unc-5(e152)* genetic background. (F) Comparing the percentage of the DTC migration defect rates observed across different experimental groups in anterior (left) and posterior (right) arms for samples shown in 4E, with more specific defect categories and classifications. Suppression of *unc-5(e152)* D/V defects was not observed in anterior arms but was evident in posterior arms. Lighter one means under RNAi control L4440 empty vector, darker one means under *lin-44* RNAi. (B-F) All sample sizes refer to individual worms. On the graphs, “no defect” means no defect observed in that group. Error bars represent the standard error of the sample proportion. Statistical analysis was performed using a pairwise proportion test, with p-values adjusted for multiple comparisons via the Benjamini-Hochberg procedure. Significant differences are indicated between groups where applicable. ****p < 0.0001 ***p < 0.001 **p < 0.01; *p < 0.05; no mark means the comparison was not statistically significant. The corresponding sample sizes and statistics are presented in Table S4.

With this double mutant, we interrogated the genetic interaction of *mom-5*/Frizzled and the *unc-5*/Netrin receptor. It is known that *mom-5* genetically represses *unc-5* (via positive regulation of Rac pathway components (Levy-Strumpf et al., 2015)). One key piece of evidence for this conclusion is that *mom-5* A/P migration defects are suppressed by *unc-5* loss of function (Levy-Strumpf et al., 2015), indicating that *mom-5* and *unc-5* affect A/P migration in opposite directions, and that *unc-5* is downstream of *mom-5* in the network. If *mig-21* functions primarily as a member of the Netrin pathway, we would expect *mig-21(u787)* either to slightly enhance the A/P defect suppression or show no effect in a *mom-5; unc-5* dual loss of function experiment. On the other hand, if *mig-21* primarily acts redundantly with Wnt pathway signaling, we might expect the overall penetrance of A/P polarity defects to increase but for these defects to still be suppressed by *unc-5* loss of function. Instead, we see something more complicated.

Otherwise, wild-type worms on *mom-5* RNAi have a ∼20-30% A/P migration defect (Fig. 2C and Fig. 4D). The *unc-5(e152)* mutant on *mom-5* RNAi shows suppression of A/P migration defects to <15% (Fig. 4B), as expected. However, the *mig-21(u787); unc-5(e152)* double mutant on *mom-5* RNAi does not suppress the A/P migration defect relative to the *mig-21(u787)* mutant on *mom-5* RNAi alone (Fig. 4D). Indeed, *mig-21(u787)* appears to synergize with *unc-5(e152)* to convert A/P migration defects to the “no turn” phenotype, which is considered to be a combination of a failed first turn and mispolarized second turn. We conclude that suppression of *mom-5* RNAi A/P migration defects by loss of *unc-5* is *mig-21*-dependent.

When we examine D/V migration defects in the same treatment groups, we see that *unc-5* loss of function causes substantial ventralization of migration, as expected, and that a substantial fraction of these is enhanced to “no turn” phenotypes by *mig-21(u787)*, and to a lesser degree by *mom-5* RNAi (Fig. 4B). Loss of function caused by *mig-21(u787)* simultaneously enhances A/P and D/V polarity defects caused by loss of function of *mom-5* and *unc-5* function, respectively.

We next tested a treatment by which loss of function of a Wnt signaling pathway member could suppress D/V migration defects caused by loss of *unc-5.* It had previously been shown that *lin-17*/Frizzled and *lin-44*/Wnt interact with *unc-5* in DTC migration (Levy-Strumpf et al., 2015). That study focused on their redundant regulation of A/P polarity, but shows data that suggest that loss of function of those Wnt pathway members suppresses the D/V defects caused by *unc-5* loss of function. When we put the *unc-5(e152)* mutant on *lin-44*/Wnt RNAi, the *unc-5(e152)* D/V defect rate is suppressed from over 80% to just over 60%. This suppression is completely eliminated in a *mig-21(u787)* mutant background, with the D/V migration defect identical to the D/V defect rate in the *mig-21(u787); unc-5(e152)* mutant on RNAi vector control (Fig. 4E-F). We thus conclude that suppression of *unc-5(e152)* D/V migration defects by loss of function of a Wnt ligand is *mig-21* dependent. Taken together, these results lead us to conclude that *mig-21* plays a key role in balancing the effects of Wnt and Netrin pathway signaling on DTC migration.

### *mig-21* is not required for cessation of DTC migration but sensitizes adult DTCs to polarizing signals

We first began investigating *mig-21* because RNA-seq data shows that it is strongly and specifically expressed in adult DTCs, however, we went on to identify larval roles for *mig-21* in the DTC. We next asked if it interacts with a regulator of the adult DTC, *vab-3*/Pax6. This transcription factor is required for the cessation of DTC migration via the transcriptional switch in the alpha integrin subtype expressed by the DTC from *ina-1* to *pat-2* (Meighan and Schwarzbauer 2007). Loss-of-function of *vab-3* causes continued DTC migration in adulthood in which the DTC takes on a meandering or curling path (Meighan and Schwarzbauer 2007). Normally, DTC migration ends at the dorsal midbody, and *mig-21(u787)* mutants also cease migration in this position (Fig. 5A). In otherwise wild-type worms with DTC markers treated with *vab-3* RNAi, we see a high penetrance of overmigrated and curling gonad tips (Fig.5A) (in excess of 90%, with ∼70% showing extra turns at the tip by 52-55 hrs post L1 arrest, Fig. 5B). RNAi knockdown of *vab-3* combined with the *mig-21(u787)* allele displays a substantial shift from continued DTC migration with extra turns to DTC overmigration along the dorsal body wall along a straight path (Fig. 5C-D). Cessation still fails, but the DTC path stays straighter. If extra turns observed after *vab-3* RNAi treatment result from chaotic DTC polarization in response to signaling gradients in the adult, we interpret the suppression of those turns by *mig-21(u787)* to reflect a role for *mig-21* in the continued sensitization of the adult DTC to Wnt and Netrin gradients, just as it is important for sensing and integrating this positional information in the larvae.

**Figure 5.**
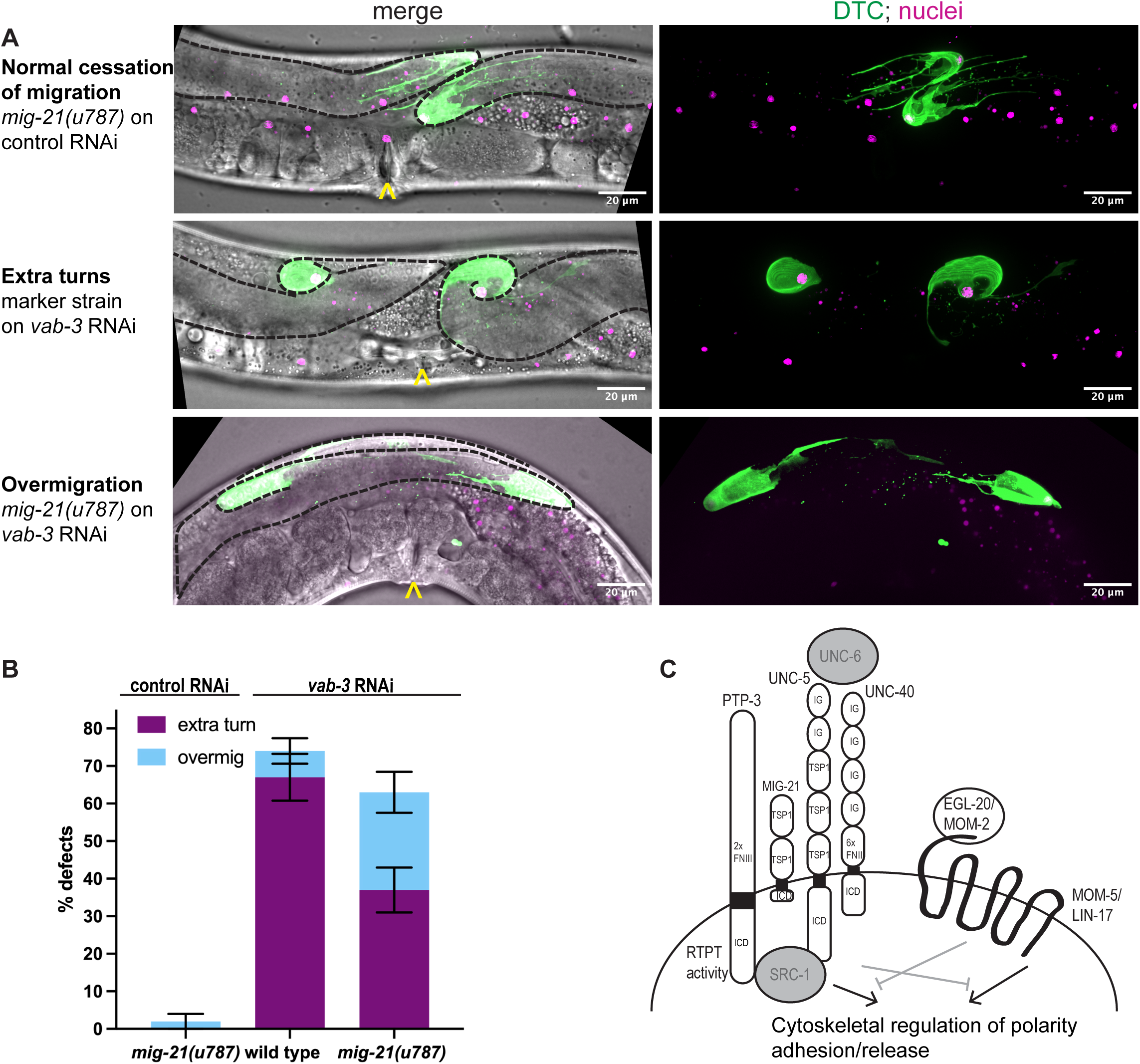
*mig-21* sensitizes adult DTCs to polarizing signals (C) Micrographs: Confocal fluorescence imaging of *C. elegans* young adult hermaphrodites expressing (*lag-2p::mNG::PH; lag-2(bmd202[lag-2::P2A::H2B::mT2]*) without (middle) and with (top and bottom) *mig-21(u787)* under control (top) and *vab-3* RNAi (middle and bottom) feeding treatment to assess DTC migration cessation defect phenotypes. Images are Z-projections through thickness of the gonad required to capture the whole distal gonad. Black dashed lines outline gonads. Yellow carats mark proximal vulval position. Scale bar, 20 μm. (B) Comparing the percentage of two main DTC migrate cessation defect rates observed across different experimental groups under *vab-3* RNAi feeding treatment. An “extra turn” to “overmigration” defect shift was observed. A robust “extra turn” defect rate marked control on *vab-3* RNAi (n=38/57) decreases with *mig-21(u787)* on *vab-3* RNAi (n=24/65), p < 0.01. However, migration cessation is not rescued; a significant increase in “overmigration” defects is observed between marked control (n=4/57) and *mig-21(u787)* on *vab-3* RNAi (n=17/65), p < 0.05. All sample sizes refer to individual worms. Error bars represent the standard error of the sample proportion. Statistical analysis was performed using a pairwise proportion test, with p-values adjusted for multiple comparisons via the Benjamini-Hochberg procedure. Significant differences are indicated between groups where applicable. ****p < 0.0001 ***p < 0.001 **p < 0.01; *p < 0.05; no mark means the comparison was not statistically significant. (C) Cartoon of MIG-21 and genetic interactors from this study, with annotations after (Norris et al. 2014). UNC-6 and SRC-1 (gray) are known to interact with UNC-5, though we have not yet examined their function in the pathway including *mig-21*.

### Conclusions and future directions

The nematode-specific *mig-21* gene encodes a thrombospondin repeat-containing protein that genetically interacts with Wnt, Netrin, and LAR RPTP receptors during cell migration in both the Q neuroblast cells and–as we have now discovered–in the DTC. In both contexts, *mig-21* loss of function more strongly affects the cell that initially migrates to the posterior (this work and Sundararajan & Lundquist, 2012), up the Wnt gradient.

The molecular basis of these interactions is not known. Previous work (Middelkoop et al. 2012) notes the thrombospondin domains shared by MIG-21 and UNC-5 could potentially mediate direct interactions between MIG-21 and UNC-40/DCC (as UNC-5 and UNC-40 were known to interact (Lim and Wadsworth 2002); the genetic evidence, in that case, supports parallel activity of the receptors (Middelkoop et al., 2012; Sundararajan & Lundquist, 2012). In the case of the DTC, our results support *mig-21* acting in the same pathway as *unc-5,* with considerable redundancy in governing ventral repulsion and A/P polarity. We note that the deletion of the C-terminal intracellular domain of UNC-5 is sufficient to abrogate UNC-5-dependent regulation of the first turn (and is known to act via *src-1*(Lee et al. 2005)), and this deficiency is not compensated for by the presence of MIG-21 (which has an intracellular domain of only 64 amino acids). Deciphering the molecular basis for UNC-5/MIG-21 redundancy in the future should therefore focus on the extracellular domains of these proteins.

MIG-21 and the UNC-40/DCC Netrin receptor are thought to regulate Wnt signal response in the Q neuroblast cells by restricting the direction of cell polarization (Sundararajan and Lundquist 2012). It has subsequently been shown that *mig-21* together with *dpy-19* regulates UNC-40 subcellular localization to the leading edge of the polarizing Q neuroblast (Ebbing et al. 2019). The proposed model by which MOM-5/Frizzled, the Wnt receptor, restricts UNC-5/Netrin receptor activity in the DTC proposed by (Levy-Strumpf & Culotti, 2014) is strikingly similar–a limitation of the direction of cell polarization. Though this has not been demonstrated molecularly yet in the DTC, in other systems, Frizzled protein itself, like UNC-40 in Q neuroblasts, is known to polarize in the cell membrane (Strutt 2001).

Our results are most consistent with a model in which MIG-21 functions in the same pathway as UNC-5, and either in parallel or redundantly with MOM-5/Frizzled in conferring A/P polarity, by which it mediates crosstalk between Wnt and Netrin signaling (Fig. 5C). Integrating MIG-21 into existing models of DTC polarization implicates several candidates worth pursuing in the future, like the non-receptor tyrosine kinase protein SRC-1 which is recruited by the intracellular domain of UNC-5 and likely leads to cytoskeletal rearrangement in support of proper polarization (Lee et al. 2005), and the ligand UNC-6/Netrin which signals through UNC-5 and UNC-40 (Norris et al. 2014) (Fig. 5C). Further work in this area will lead to the integration of known regulators and novel candidates into a more complete model of DTC migration.

## Methods

*Sections of this text are adapted K. Gordon lab publications (Li et al. 2022; Singh et al. 2024) as they describe our standard laboratory practices*.

### Target gene selection

The target gene *mig-21* was selected from the single-cell transcriptional atlas of young adult *C. elegans* WormSeq.org app (https://wormseq.org) designed by Ghaddar et al. (2023). The app offers various tools for gene expression analysis, including but not limited to identifying specific gene markers and assessing gene expression across cell types. In our study, we used the “top gene markers” function first to view the top 100 gene markers for the distal tip cell. This list was generated by the Monocle3 (https://cole-trapnell-lab.github.io/monocle3/) “find markers” function, which “measures specificity using the Jensen-Shannon distance”. We first sorted the candidates by the highest “marker score” which yields genes that are both relatively specific and highly expressed, then we also sorted them by the “specificity” score as recommended by https://wormseq.org. We found that *mig-21* had the highest marker score and ranked third in specificity, and was the only gene present in the top ten of both sorts (Figure 1A).

### Strains

Some strains were provided by the CGC, which is funded by NIH Office of Research Infrastructure Programs (P40 OD010440). In strain descriptions, we designate linkage to a promoter with a p following the gene name and designate promoter fusions and in-frame fusions with a double semicolon (::). Some integrated strains (xxIs designation) may still contain for example the *unc-119* (ed4) mutation and/or the *unc-119* rescue transgene in their genetic background, but these are not listed in the strain description for the sake of concision, nor are most transgene 3’ UTR sequences. The wild-type strain N2 was used as a control. New strains produced for this study are: KLG041 *cpIs122[lag-2p::mNeonGreen:: PLC^δPH^] II; mig-21(u787) III; lag-2(bmd202[lag-2::P2A::H2B::mT2] ^lox511I^2xHA) V,* and KLG042 *mig-21(u787) III; unc-5(e152) IV*.

### Worm rearing

*C. elegans* strains were maintained at 20°C on standard NGM media and fed *E. coli* OP50 for routine strain maintenance. All animals assessed were hermaphrodites, as males have nonmigratory DTCs. Worm populations were synchronized at L1 arrest for developmental staging by standard egg preps (Stiernagle 2006).

### Confocal imaging

All images were acquired on a Leica DMI8 with an xLIGHT V3 confocal spinning disk head (89 North) with a ×63 Plan-Apochromat (1.4 NA) objective and an ORCAFusion GenIII sCMOS camera (Hamamatsu Photonics) controlled by microManager (Edelstein et al. 2010). mNG was excited with a 488 nm laser, and mT2 was excited by a 445 nm laser. Worms were mounted on 4% noble agar pads in 0.01 M sodium azide (VWR (Avantor) Catalog Number 97064-646) for live imaging.

### RNAi

*E. coli* HT115(DE3) containing the L4440 plasmid, either with or without a dsRNA trigger insert sourced from the Ahringer or Vidal Unique RNAi libraries, or our own clone in the case of *unc-5* (Singh et al. 2024), were cultured overnight from single colonies at 37°C with ampicillin (100 μg/mL, VWR (Avantor), Catalog no. 76204-346). Subsequently, dsRNA expression was induced with 1mM IPTG (Apex BioResearch Products, cat # 20-109) for one hour at 37°C, followed by plating 200 μl of the culture and incubating overnight at room temperature on prepared NGM plates with a 1:1 ratio (2.5 μL each) ampicillin and IPTG spread uniformly on the surface with a glass spreader. Worm populations were synchronized by bleaching according to a standard egg prep protocol (Stiernagle 2006), plated on NGM plates seeded with RNAi-expressing bacteria as arrested L1 larvae, and kept on RNAi until the time of imaging. RNAi treatment was conducted at 20°C.

### Image analysis

Images were processed in FIJI (Version: 2.14.1/1.54f) (Schindelin et al., 2012). Detailed descriptions of image analysis for different experiments are provided below.

### Measurements of DTC length and DTC nuclear location

The DTCs were identified in the fluorescence images (Fig. 1E-F). The length of the DTC was determined as the distance from the gonad tip to the farthest point of the DTC edge. The location of the DTC nucleus was defined as the distance from the anatomical gonad tip to the center of the DTC nucleus. All measurements were obtained using the FIJI straight line tool.

### Staging and scoring of DTC migration defects

L4 (Figures 1-4, Supplement) or young adult (Figure 5) animals were scored for DTC migration defects based on gonad morphology. In Figures 1-4, we used the framework of (Levy-Strumpf & Culotti, 2014) to categorize defects of A/P polarity, D/V polarity, and “no turn” defects in which D/V turning fails and A/P polarity is reversed. Some cases of A/P polarity defects result in DTC migration into the pharynx (anterior) or tail (posterior) regions, and others involve extra turns in which migration on the dorsal body wall started in these directions and subsequently reversed back towards the midbody. To these we also add a “severe” category in which gonad outgrowth fails completely or the gonad forms a disorganized mass, and an “other” category, usually cases in which the last phase of gonad migration is not maintained along the dorsal body wall. In Figure 5, *vab-3* RNAi causes failure of migration cessation and perpetual migration, and we separate specimens into classic *vab-3* phenotypes in which the DTC makes extra turns and cases of “overmigration” in which the DTC maintains its path along the dorsal body wall but overshoots the midbody.

### Quantification and statistical analysis

Statistical tests, sample sizes, test statistics, and p-values for each analysis are provided in the corresponding figure legends. Statistical analysis and multiple comparison corrections were performed using GraphPad Prism Version 10.4.0 (527) for macOS (GraphPad Software, Boston, MA, USA), and Rstudio version 2024.12.0+467 with the rstatix package (Kassambara 2023). The pairwise proportion test was used to compare proportions between experimental groups, and p-values were adjusted for multiple comparisons using the Benjamini-Hochberg procedure to control the False Discovery Rate (FDR) at 0.05. For histograms presented in the Figures, the standard error of the sample proportion was calculated with the *SEp^^^ = √p^^^(1 − p^^^)/n*, where *p^^^* is the proportion of specimens (worms or gonads) of the total observed *(n)* with the phenotype; error bars reflecting these standard errors (expressed as percentages) were added to each plot using GraphPad Prism. Unpaired Student’s two-tailed t-test was also performed to compare the means of DTC length and DTC nuclear location measurements, with p-values also calculated with GraphPad Prism.

## Acknowledgments

We would like to thank members of the Gordon Lab, especially Noor Singh and Camille Miller, for their technical help and feedback. We thank Rob Dowen and Peter Breen for sharing RNAi clones. Some strains were provided by the CGC, which is funded by NIH Office of Research Infrastructure Programs (P40 OD010440). Funded by NIGMS Grant R35GM147704 to KLG.

## Author Contributions

X.L. and K.L.G. conceived the project and designed the experiments. X.L. carried out the experiments. X.L. and K.L.G. analyzed the data and wrote the manuscript.

